# Polygenic Associations between Motor Behaviour, Neuromotor Traits, and Active Music Engagement in Four Cohorts

**DOI:** 10.1101/2025.03.27.645667

**Authors:** T.L. Henechowicz, P.L. Coleman, D.E. Gustavson, Y.N. Mekki, S. Nayak, R. Nitin, A.C. Scartozzi, E.S. Tio, R van Klei, D. Felsky, M.H. Thaut, R.L. Gordon

## Abstract

Phenotypic investigations have shown that actively engaging with music, i.e., playing a musical instrument or singing may be protective of motor decline in aging. For example, music training associated with enhanced sensorimotor skills accompanied by changes in brain structure and function. Although it is possible that the benefits of active music engagement “transfer” to benefits in the motor domain, it is also possible that the genetic architecture of motor behaviour and the motor system structure may influence active music engagement. This study investigated whether polygenic scores (PGS) for five behavioural motor traits, 12 neuromotor structural brain traits, and seven rates of change in brain structure traits trained from existing discovery genome-wide association studies (GWAS) predict active music engagement outcomes in four independent cohorts of unrelated individuals of European ancestry: the Canadian Longitudinal Study on Aging (CLSA; N=22,198), Wisconsin Longitudinal Study (WLS; N=4,605), Vanderbilt’s BioVU Repository (BioVU; N=6,150), and Vanderbilt’s Online Musicality study (OM; N=1,559). Results were meta-analyzed for each PGS main effect across outcomes and cohorts, revealing that PGS for a faster walking pace was associated with higher amounts of active music engagement. Within CLSA, a higher PGS for walking pace was associated with greater odds of engaging with music. Findings suggest a shared genetic architecture between motor function and active music engagement. Future intervention-based research should consider the genetic underpinnings of motor behavior when evaluating the effects of music engagement on motor function across the lifespan.

## 1. Introduction

Actively engaging with music, i.e., playing a musical instrument or singing, is an underappreciated lifestyle factor in healthy ageing because it is a cognitively stimulating, involves motor learning, coordination, and multisensory integration (1). Research suggests that skills gained from playing instruments or singing may “transfer” to enhanced skills in a non-related domain (e.g., cognitive or motor skills) and changes in brain structure and function (1–3). Motor function is an integral part of neurocognitive health in aging, where motor decline precedes cognitive decline (4,5). There are several lines of phenotypic evidence showing how active music engagement may be protective of motor function. For example, a neuroimaging meta-analysis showed that musicians, compared to non-musicians, have greater auditory-motor network connectivity and structural adaptations in regions important for motor control including the corpus callosum, internal capsule, and sensorimotor and subcortical areas (6). Behaviourally, musicians, compared to non-musicians, have enhanced musical motor skills (e.g., audio-motor synchronization) (7–9), better performance on standardized motor function assessments (10,11), faster reaction time in spatial (12), multisensory integration (13), and visuomotor tasks (14), and enhanced motor sequence learning and retention (15,16). Music training interventions may also heighten motor skill development in children and adolescents (17,18). Given what is now known about genetic contributions to brain development and plasticity (19–21), it is possible that these “transfer” effects are partially influenced by shared genetic variation (22). The current investigation seeks to understand how genetic factors for motor behaviour and neuromotor traits may influence individual differences in music engagement.

Given the clear behavioural and neural links between musical instrument training and the motor system, several authors hypothesize that long-term or regular playing of an instrument or singing is likely an environmental factor that modulates motor system neural plasticity (23). In parallel, prior individual differences in motor skills or neuromotor features may contribute to who seeks out musical training (2,3,24), and prior work does not rule out the possibility that alternative or additional shared biology, i.e., shared genetic architecture, underlies these phenotypic correlations. For example, the potential “transfer” of childhood musical instrument engagement to verbal ability four years later in adolescence is in part driven by shared genetic effects (25). To investigate the hypothesized shared genetic architecture between active music engagement and motor phenotypes, we can adapt insights from a recent musicality cross-trait framework, the *Musical Abilities, Pleiotropy, Language, and Environment* (MAPLE) (22). The MAPLE framework makes a case that both musical abilities and musical environments may arise from genetic effects, i.e., the niche-picking phenomenon where individuals are drawn to environmental influences for which they have higher genetic liability to well adapted. Second, commonly found phenotypic associations between musicality and motor traits can be explained through partially shared genetic architecture and mediating neurobiological phenotypes (22).

Indeed, active music engagement and motor traits are moderate to highly heritable. Music-related traits have an average heritability of 42% (26), including 78% for musical instrument engagement (25), 33–66% for musical aptitude (27), and 41–69% for musical practice (28,29). Recent genomic studies show evidence for SNP-based heritability of 12% for music engagement (30). For motor traits, a twin study found evidence for 68% and 70% heritability of motor control and motor learning, respectively (31). Additionally, genetic studies of neurobiological endophenotypes, e.g., the thickness, volume, and change in volume/thickness of motor system brain structures, are available publicly through *Enhancing Neuroimaging Genetics through Meta-analysis Consortiums’* GWASs and show evidence for the heritability and detectable polygenic signals (19,32,33). Accordingly, cross-trait population-level genomic methods, i.e., polygenic scores, can leverage existing GWASs for behavioural motor and neuromotor phenotypes to understand their shared biological etiology with active music engagement.

In this pre-registered study (https://doi.org/10.17605/OSF.IO/SQ2NC), we examined whether PGSs for three categories of motor traits—behavioural, brain structure, and rate of change of brain structure—predict active music engagement in four cohorts: the Canadian Longitudinal Study on Aging (CLSA) (34,35), Vanderbilt’s BioVU repository (BioVU) (36,37), Wisconsin Longitudinal Study (WLS) (38), and the Vanderbilt’s Online Musicality Study (OM) (30). We predicted that greater genetic predisposition for motor function (i.e., greater PGS for walking pace, lower PGS for reaction time, lower PGS for hand muscle weakness, lower PGS for clinical motor diagnosis of Parkinson’s disease, and lower PGS for Developmental Coordination Disorder risk) would each be associated with higher amounts of music engagement. For neuromotor phenotypes, we predicted associations between genetic predispositions for brain structure and its rates of change in brain structure with active music engagement, without specific directional hypotheses.

## 2. Methods

### 2.1. Data

#### Study populations (target cohorts)

Four large cohort studies were analyzed: CLSA (N=22,198), WLS (N=4,605), BioVU (N=6,150), and OM (N=1,559). Given the limited data availability in these cohorts, analyses were constrained to individuals of European ancestry (see **Supplementary Methods 1.1-1.4** for phenotyping and quality control protocols). We extracted active music engagement phenotypes and categorized the outcomes into constructs including music practice, music achievement, and music engagement (see **Supplementary Table 1.** and **Supplementary Methods 1.1-1.4**).

#### Discovery GWAS summary statistics used for PGS construction

PGSs were constructed using summary statistics from published GWASs of motor behaviour traits (5 traits), structural motor brain traits (12 traits), and the rate of change of brain structure across the lifespan (19) (7 traits) (see **Appendix B. Supplementary Table 2.** for descriptions of the discovery GWAS). All discovery GWAS samples were non-overlapping with target cohorts and were of European ancestry to match the target cohorts’ ancestries to avoid population stratification issues(39).

#### Motor behaviour discovery GWAS

We selected motor behaviour traits from the GWAS atlas (https://atlas.ctglab.nl/), catalogue (https://www.ebi.ac.uk/gwas/), and from a general literature search. Available GWASs included reaction time(40), self-reported walking pace (41), hand muscle weakness (grip strength <30 kg in males and <20 kg in females). (42), motor coordination difficulties in children (43), and Parkinson’s disease case status (44) (see **Appendix B. Supplementary Table 2.**).

#### Neuromotor brain structure discovery GWAS

We selected recent neuroimaging GWASs from the ENIGMA consortium for PGS of structural brain traits (32,33). For subcortical regions, we selected four of seven structures (32), the nucleus accumbens, pallidum, putamen, and caudate, because of their importance for motor learning and music training (6,45). For cortical structures, we extracted Grasby et al.’s meta-GWASs (33), specifically, we selected overall cortical thickness and six cortical thickness phenotypes from regions essential to motor and sensory systems: the precentral, postcentral, inferior parietal, insula, middle temporal, and superior temporal gyri. We also included Tissink et al.’s GWAS of combined cerebellar grey matter and cerebellar white matter volume (46).

#### Neuromotor rate of change discovery GWAS

Lastly, we included Brouwer et al.’s meta-GWAS of change rates in brain structure (19). We selected a subset of brain regions and phenotypes that matched our cross-sectional analyses and had SNP-based heritability greater than 0, i.e., the total brain volume, total cerebellar white matter volume, total cortical volume, total cortical thickness, pallidum volume, putamen volume, and nucleus accumbens volume. Rate-of-change PGSs were calculated using meta-GWAS summary statistics, which meta-analyzed SNP effects independent of age differences between cohorts. Therefore, positive PGSs represent the genetic liability for positive change (more growth or less shrinkage) across the lifespan, and negative PGSs represent negative change (less growth or more shrinkage) across the lifespan.

#### PGS construction

PGSs were calculated using PRS-CS or PRS-CS-auto, which use a Bayesian regression framework and place a continuous shrinkage prior on SNP effect sizes (47), outperforming traditional clumping and thresholding methods (48). We used PRS-CS-auto to construct PGSs for the GWAS summary statistics with N>200,000 and PRS-CS with phi=0.01 for smaller sample sizes (i.e., neuroimaging and motor coordination GWASs) as suggested for highly polygenic traits (47). All PGSs were calculated with default parameters, a=1 and b=0.5 (47), and with the 1000 Genomes Project Phase 3 European linkage disequilibrium (LD) reference panel (47,49).

### 2.2. Statistical Analysis

Statistical analyses were performed using R statistical software v4.0.2. Linear models were used to test for the main effects of each motor PGS on all music engagement outcomes within each cohort. For each outcome, 24 models were fit, each including one of the 24 motor PGSs with covariates of 10 genomic principal components, age, and biological sex, yielding 168 models. Continuous and binary outcomes were modelled with ordinary least squares linear and logistic regression, respectively. As pre-registered, multiple testing corrections were applied to PGS main effect *p*-values for the total number of models within each cohort (e.g., OM has 4 outcomes × 24 models for 72 tests; WLS has 2 outcomes × 24 models for 48 tests; BioVU has 1 outcome × 24 models for 24 tests; CLSA has 1 outcome × 24 models for 24 tests). Multiple test correction was assessed using the Benjamini-Hochberg false discovery rate (FDR) procedure, with a corrected *p*-value (*q*FDR) of *q*FDR<0.05 indicating a significant effect. We assessed model fit indices of *R^2^* for continuous outcomes and Nagelkerke-Pseudo-*R^2^* and *AUC* for binary outcomes. To evaluate out-of-sample performance and overfitting, we performed resampling analyses using 100 iterations of the .632 bootstrapping method commonly applied in GWAS (50,51). In secondary analyses, we fitted the same 168 models with PGS-by-sex interactions, applying the same multiple-test correction approach. Influential observations were identified using Cook’s distance (threshold of > 4/N-*k*-1), and we conducted sensitivity analyses by removing influential observations.

Lastly, we conducted meta-analyses of the effect sizes for each PGS association across cohorts. Since outcomes were continuous and dichotomous, we converted log odds ratios to standard mean differences prior to meta-analysis (52). Inverse-variance weighted random mixed-effect meta-analyses were conducted using the restricted maximum likelihood estimator with 150 maximum iterations from the *metafor* package in R (53). We applied FDR multiple testing correction for the 24 meta-analyses.

## 3. Results

### 3.1. Descriptive Statistics

**Table 1** has descriptive statistics for N=22,198 unrelated individuals from CLSA (62.99±10.15 years old, 50% female, 77.55% had completed a post-secondary degree/diploma), N=4,605 from WLS (64.22±2.50 years old, 51% female, and the average education was 13.89±2.40 years), N=6150 from BioVU (53.13±16.38 years old, 41% female, mean Area Deprivation Index of 0.33±0.12), and N=1559 from OM (45.85±16.33 years old, 74% female, 79.74% had at least a bachelor’s degree or equivalent). Distributions of the continuous outcomes in OM are in **Figure 1**. For case-control cohorts, we compared the available socio-economic and education variables (See Note in **Table 1**). See **Supplementary Methods 1.1-1.4.** for inclusion/exclusion criteria and **Figures S1A–S1D** for correlations between age, sex, and PGSs within each cohort.

**Table 1.**
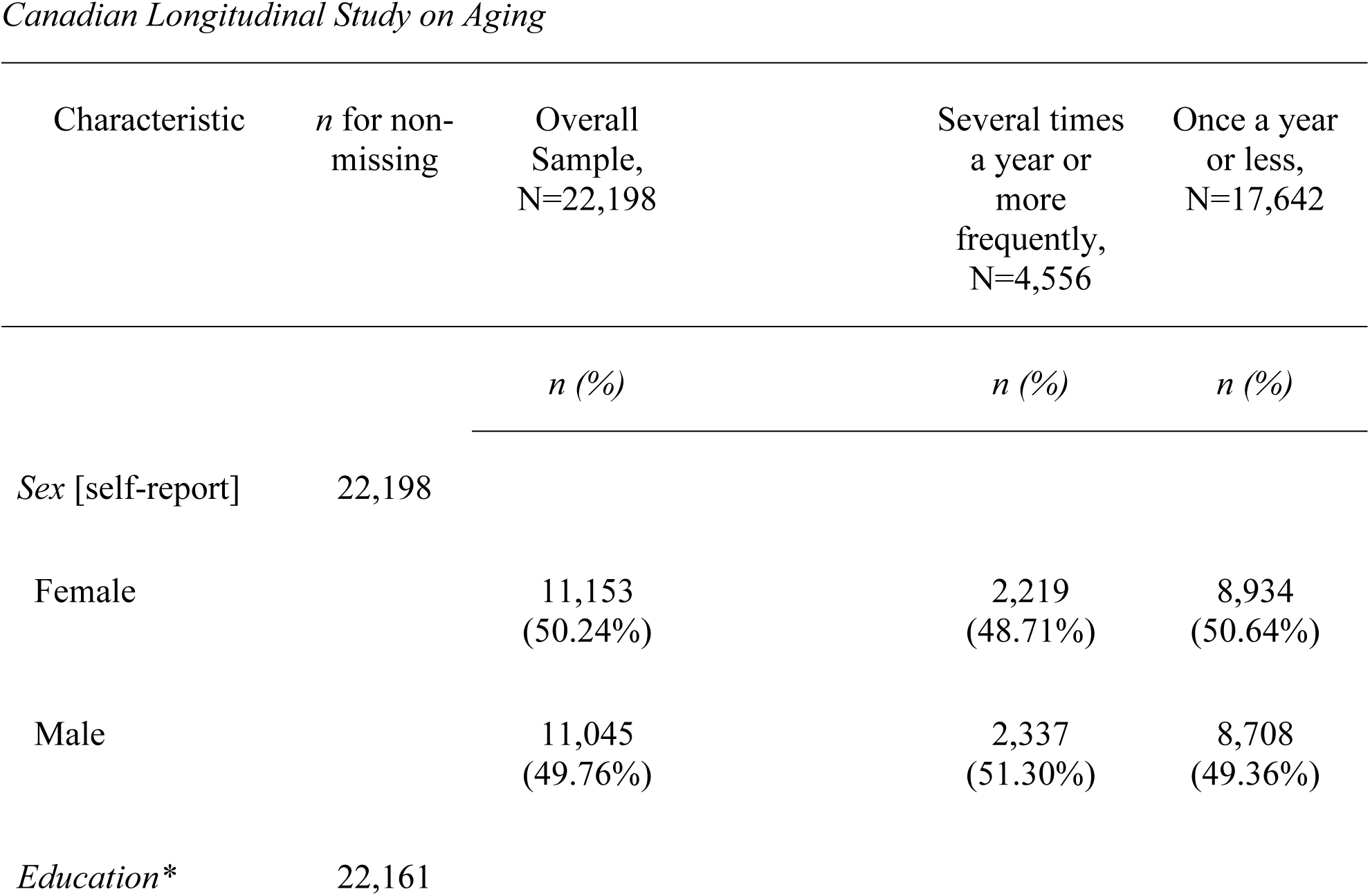
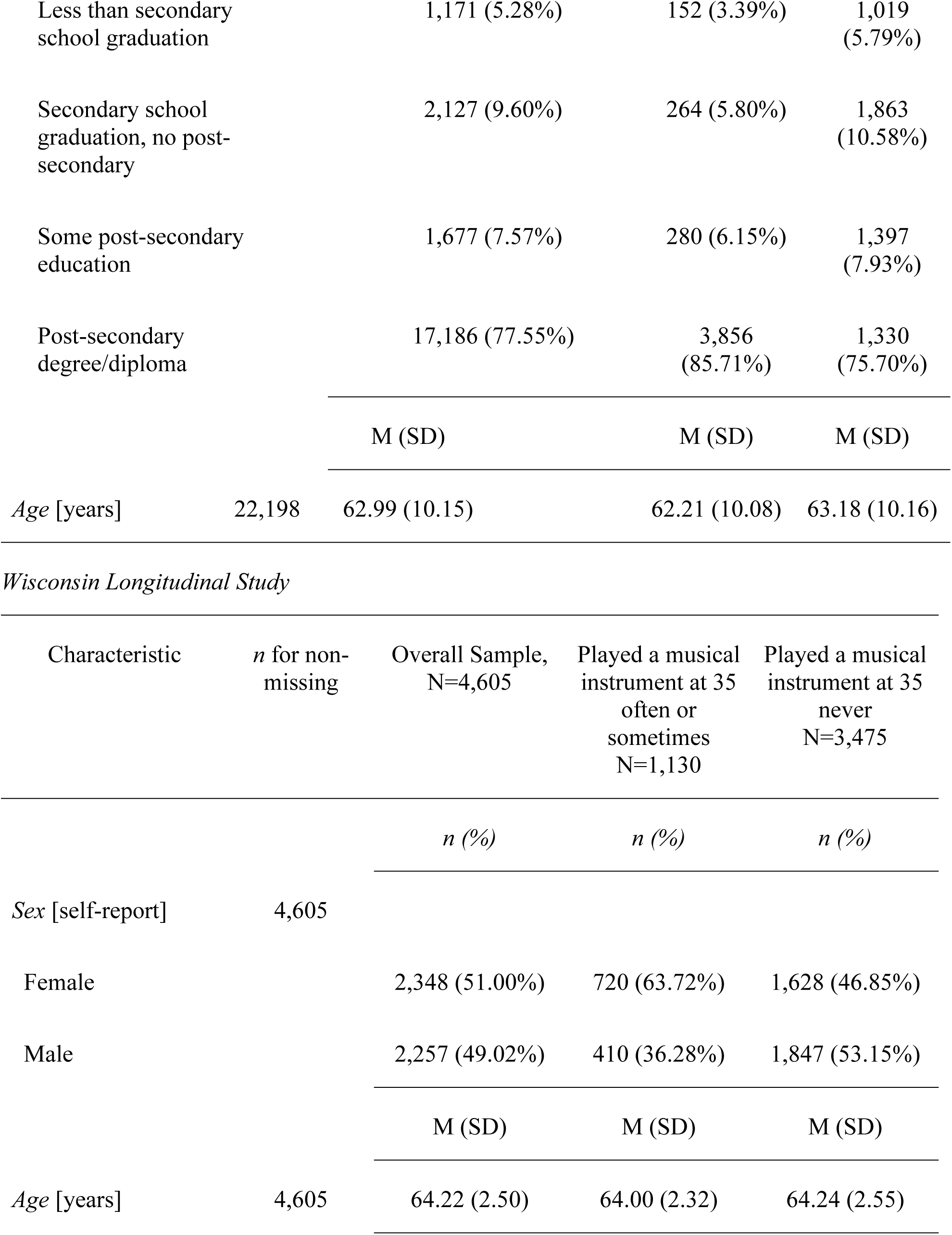
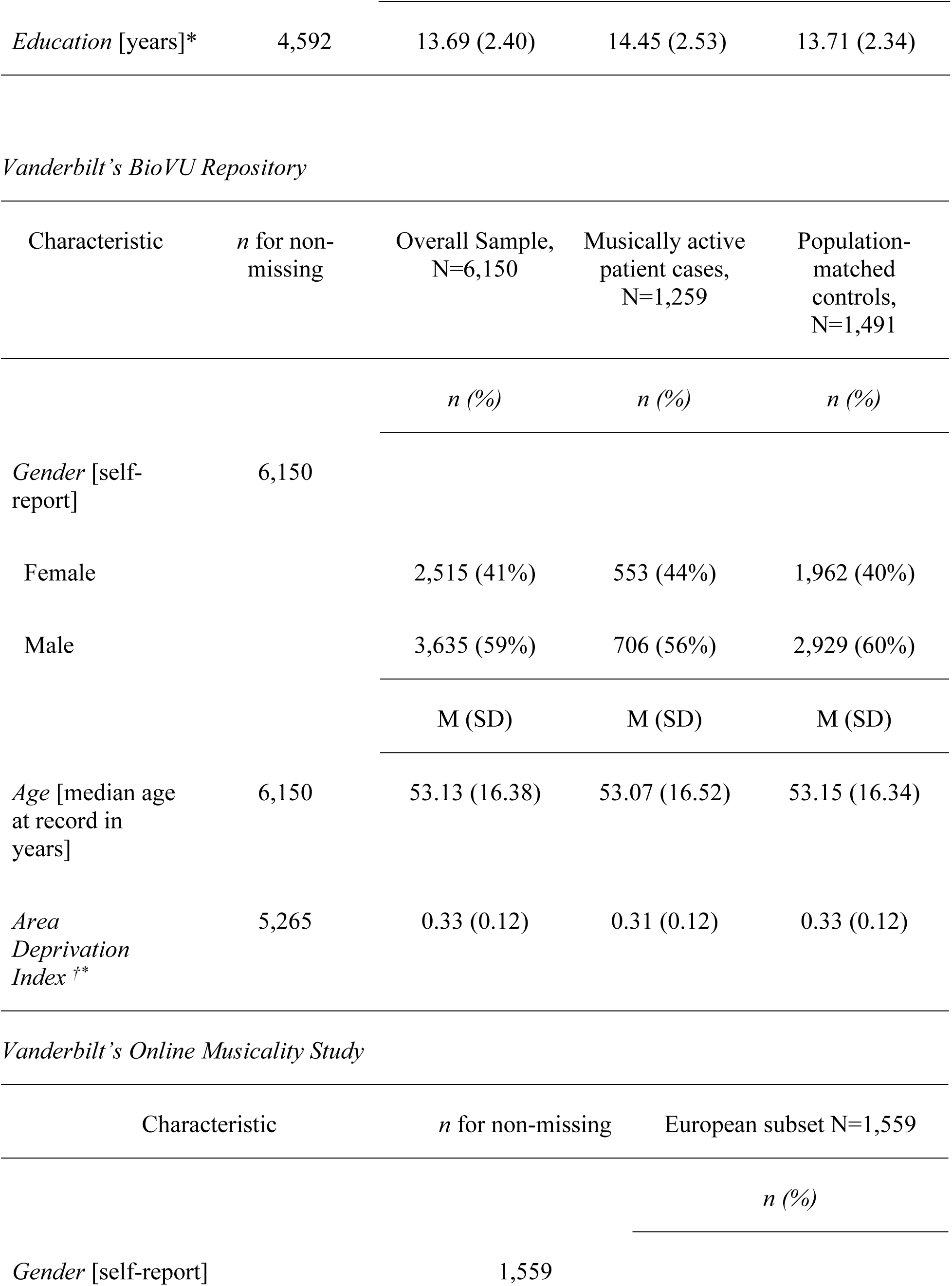
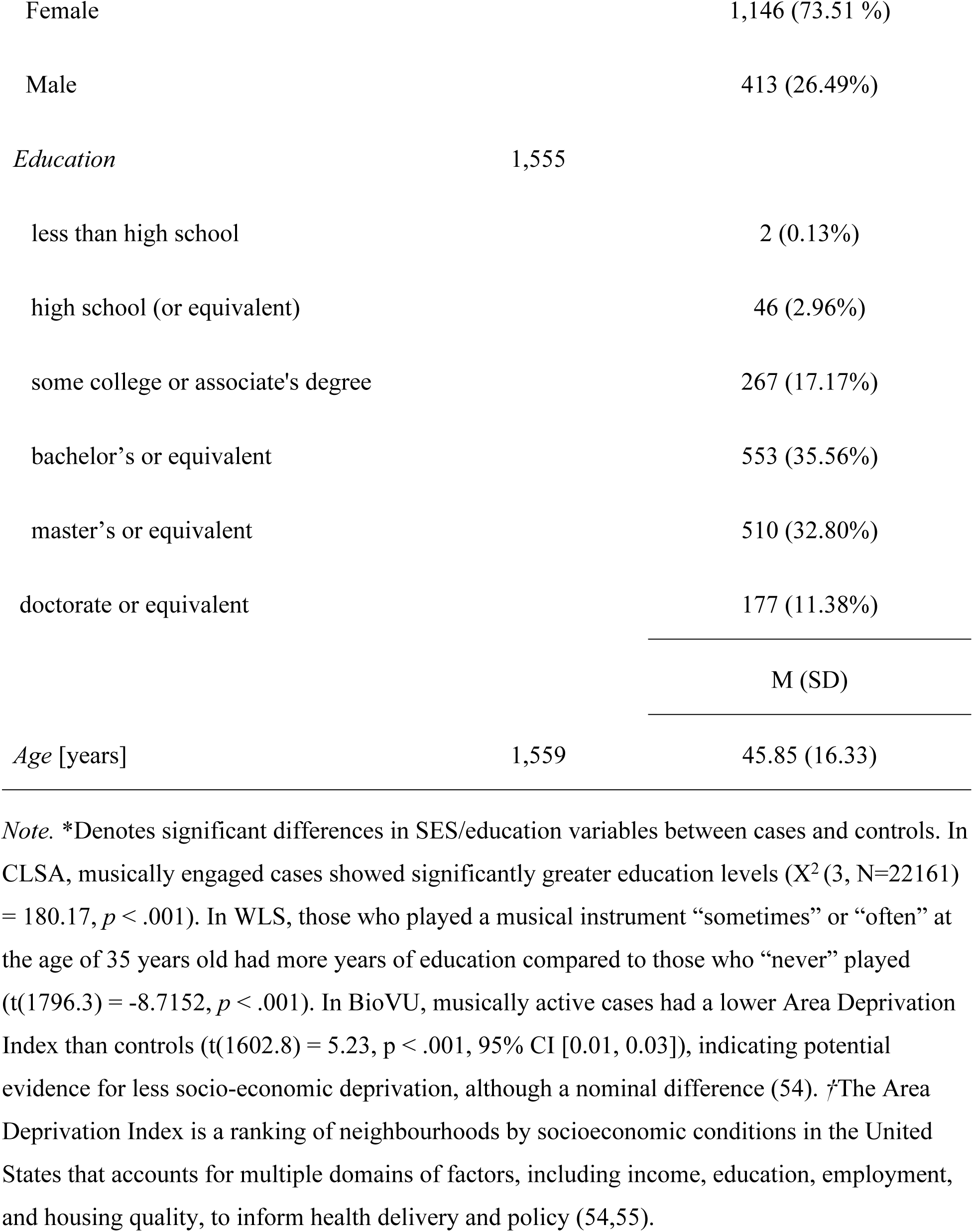
Descriptive Statistics for Target Cohorts.

**Figure 1.**
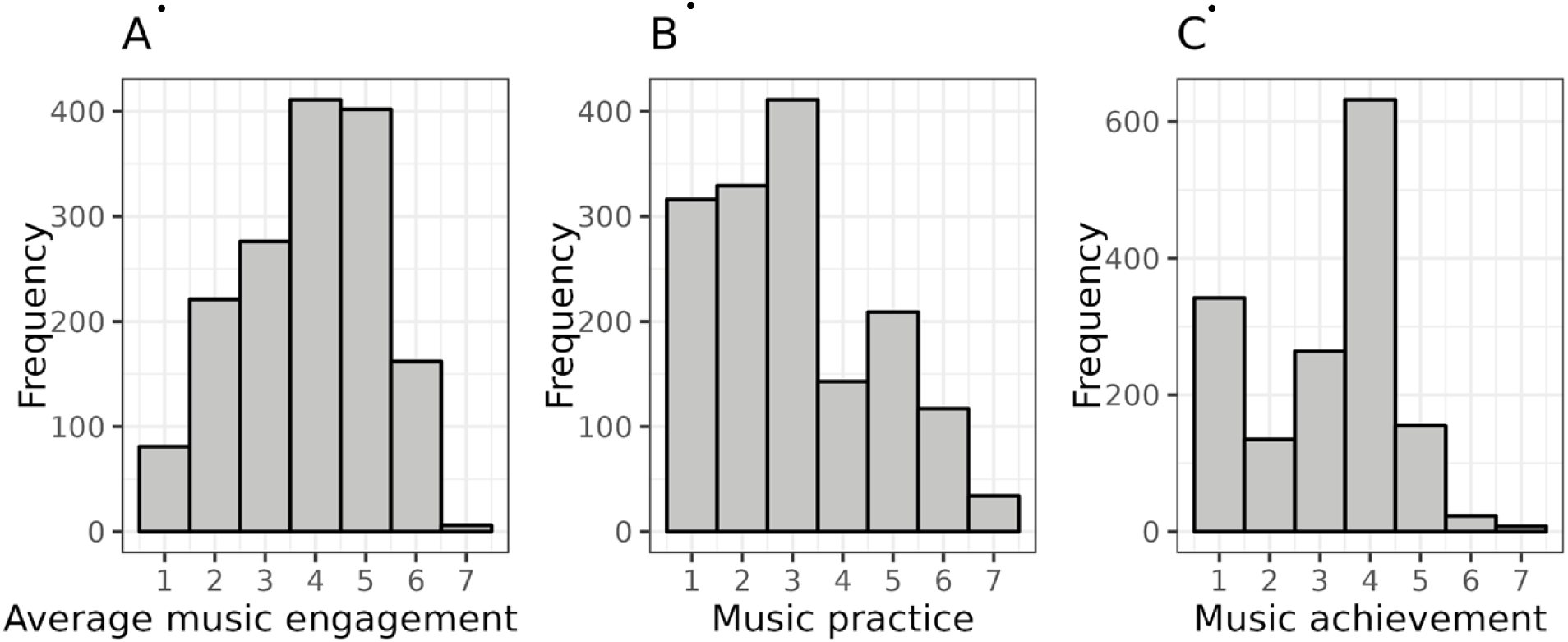
Distributions of Continuous Outcomes from Vanderbilt’s Online Musicality Study (OM). *Note.* (A) the distribution of average music engagement score, (B) music practice (as defined by question 33 in the gold-MSI), and music achievement (as defined by the creative achievement questionnaire). See **Supplementary Methods 1.4** for further phenotyping definitions in the OM cohort. Participants’ mean average score for music engagement was 3.99±1.36, and the median was 4.25 (IQR=3.25,5.25). The median self-report music practice hours was 3, corresponding to 1 hour per day (IQR=2,4). The median of self-report music achievement was 4, corresponding to “I have played or sung, or my music has been played in public concerts in my hometown, but I have not been paid for this” (IQR=3,5). All three music engagement phenotypes were significantly positively correlated: *r_s_*=0.75 (*p*<0.001) for average music engagement and music practice, *r*_s=_ 0.84 (*p*<0.001) for average music engagement and music achievement, and *r*_s_=0.58 ( *p*<0.001) for music achievement and music practice.

### 3.2. Meta-analyses of PGS Models

The meta-analyzed PGS for walking pace was positively associated with active music engagement (*b=*0.04, 95% CI [0.03, 0.05], *p*<0.001, *q*FDR<0.001) with no significant heterogeneity among studies (*Q*(6)=1.68, *p*=0.95). At an uncorrected threshold of *p*<0.05, PGS for reaction time was negatively associated with active music engagement (*b*=-0.02, 95% CI [-0.03, -0.005], *p*=0.008, *q*FDR=0.09) with no significant heterogeneity among studies (*Q*(6)=7.10, *p*=0.31) and PGS for hand muscle weakness was negatively associated with active music engagement (*b*=-0.02, 95% CI [-0.04, -0.003], *p*=0.02, *q*FDR=0.14) with no significant heterogeneity among studies (*Q*(6)=4.46, *p*=0.61) (See **Figure 2A**). Additionally, PGS for inferior parietal gyrus thickness was positively associated with active music engagement (*b=*0.02, 95% CI [0.003, 0.04], *p*=0.02, *q*FDR=0.14) with no significant heterogeneity among studies (*Q*(6)=9.44, *p*=0.15) (See **Figure 2B**).

**Figure 2.**
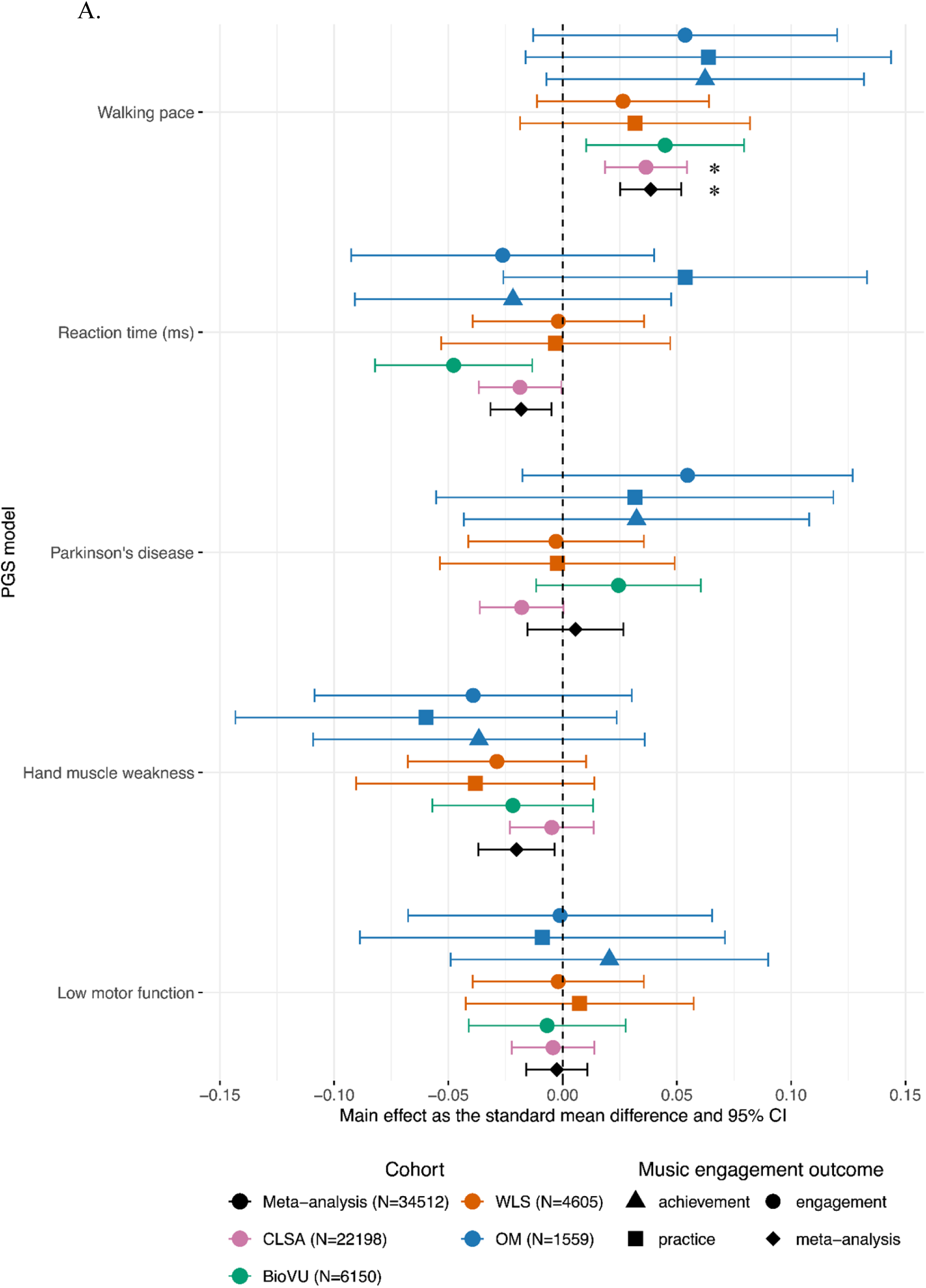
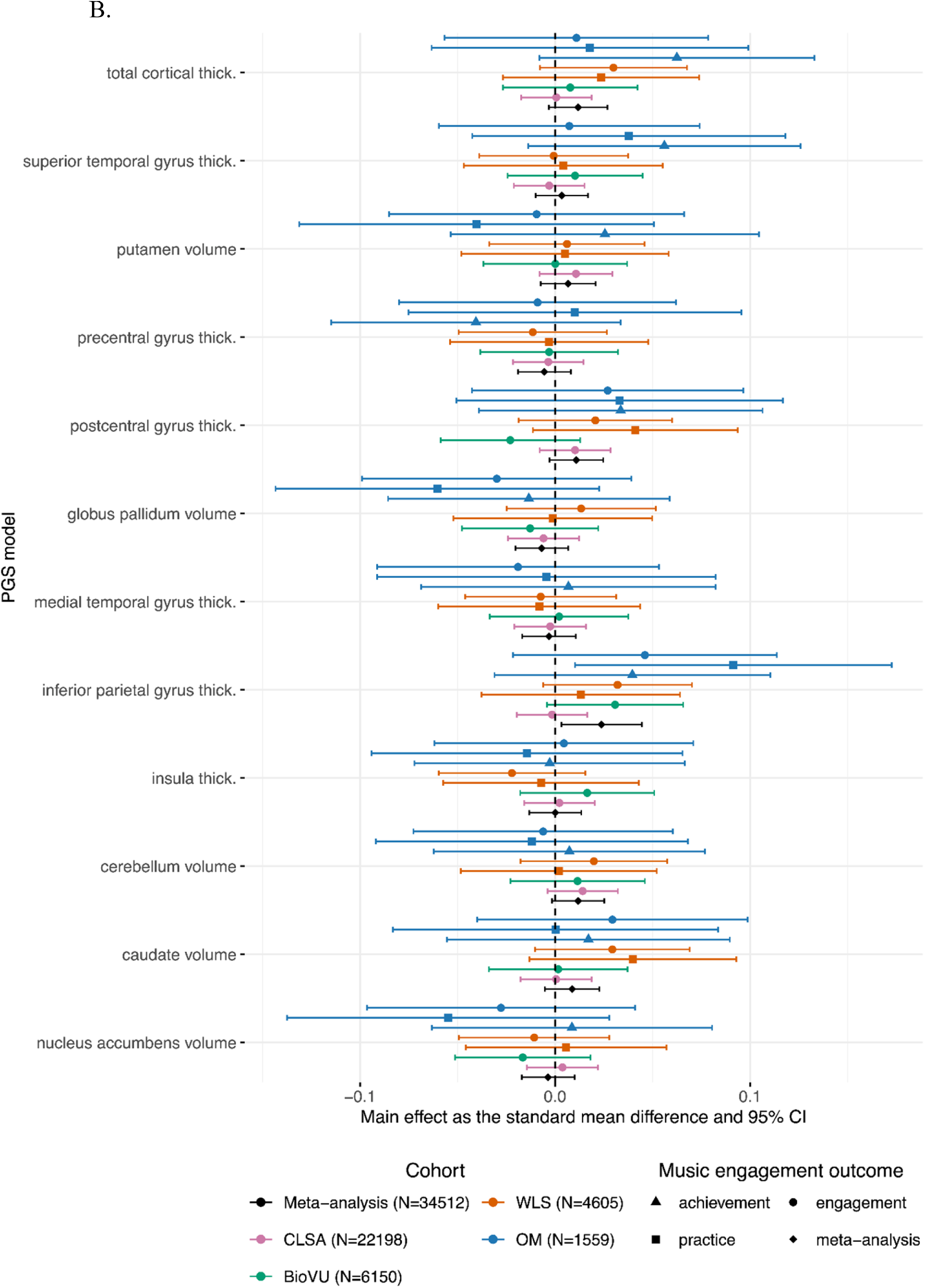
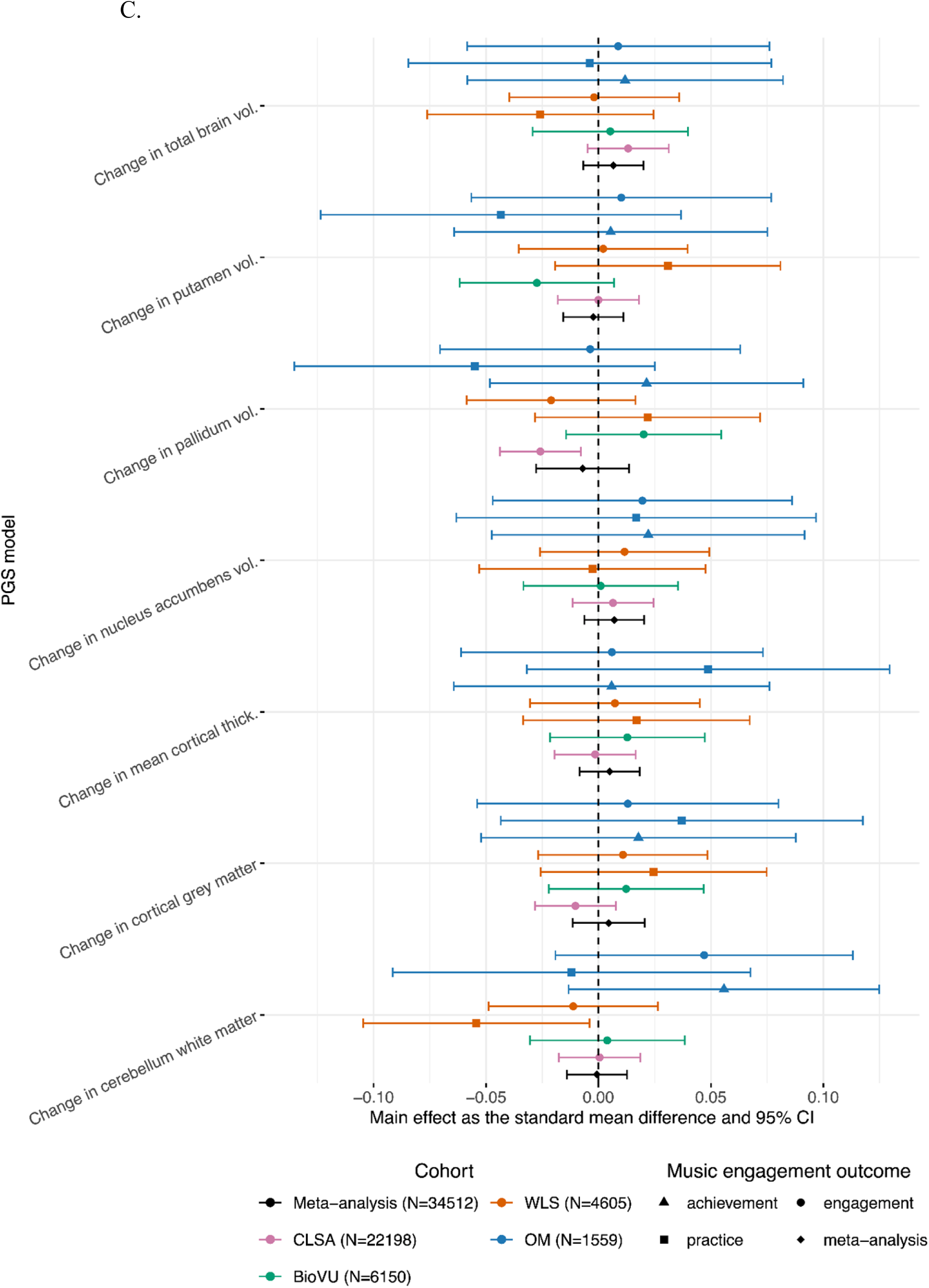
Forest Plot of 24 PGS Main Effects and Meta-analyses Across Outcomes

### 3.3. Single PGS Model Results

#### Main Effects of Motor Behaviour PGSs on Music Engagement Outcomes

PGS for walking pace was positively associated with greater odds of music engagement only in CLSA (OR =1.07, 95% CI [1.03, 1.10], *p*<0.001, *q*FDR=0.002) (see **Figure 2A**). At an uncorrected threshold of *p*<0.05, PGS for walking pace was associated with greater odds of music engagement in BioVU (OR=1.08, 95% CI [1.02, 1.15], *p*=0.01, *q*FDR=0.13). At an uncorrected threshold of *p*<0.05, there was suggestive evidence that higher PGS for reaction time (i.e., slower reaction time) was associated with lower odds of music engagement in CLSA (OR=0.97, 95% CI [0.94, 1.00], *p*=0.04, *q*FDR=0.33) and in BioVU (OR=0.92, 95% CI [0.86, 0.98], *p*=0.007, *q*FDR=0.13). See **Supplementary Table 3** for all main effects and boot-strapped performance indices and **Supplementary Table 4** for sensitivity analysis results.

#### Main Effects of Structural Brain PGSs on Music Engagement Outcomes

At an uncorrected threshold of *p*<0.05, PGS for inferior parietal gyrus thickness was positively associated with music practice in OM (*b=*0.09, 95% CI [0.01, 0.17], *p*=0.03, *q*FDR=0.93) (See **Figure 2B**).

#### Main effects of Neuromotor Rate of Change PGSs on Music Engagement Outcomes

At an uncorrected threshold of *p*<0.05, there were potential associations between neuromotor rate of change PGSs and music engagement, although there was no consistency across cohorts (See **Figure 2C**). In CLSA, PGS for the rate of change in pallidum volume was negatively associated with odds of being a musically active case (OR=0.95, 95% CI [0.92, 0.99], *p=*0.005, *q*FDR=0.06), suggesting that engaging with music is associated with a genetic predisposition for more shrinkage of the pallidum. In WLS, PGS for the rate of change in cerebellum white matter volume was associated with decreased odds of practicing a musical instrument (OR=0.91, 95% CI [0.83, 0.99], *p=*0.03, *q*FDR=0.90), i.e., greater likelihood of music practice was associated with a higher genetic predisposition for more shrinkage of the cerebellum white matter.

#### Interaction Effects of Sex on PGS Models

There were no significant sex-by-PGS interactions after multiple testing corrections. However, at the suggestive *p<*0.05 threshold, there were some potential interaction effects of sex and PGS for walking pace (See **Supplementary Table 5** and **Figures S2–9**).

All effects represented in a forest plot as effect sizes and 95% confidence intervals. Meta-analyzed effects are represented as diamonds. For continuous outcomes, main effects were estimated using linear regression and are represented as an effect size. For case-control or dichotomized outcomes, logistic regression models were used to calculate log odds, which were then transformed into standard mean differences. The results are panelled by PGS category (a) motor behaviour (5 PGSs), (b) neuromotor brain structure (12 PGSs), and (c) neuromotor rate of change in regional volume/thicknesses (7 PGSs). Asterisks (*) denote significant meta-effects after multiple-test correction (*q*FDR*<*0.05). See **Supplementary Table 1.** for cohort information and **Supplementary Table 2.** for GWAS discovery source studies. **Supplementary Table 6.** has source data for plotting.

## 4. Discussion

Our investigation revealed insights into the potential shared genetic relationship between music engagement and motor traits. First, the genetic predisposition for enhanced motor function, i.e., higher PGS for walking pace, was significantly associated with greater music engagement in CLSA and once meta-analyzed across four cohorts. We also observed potential associations between greater PGSs for inferior parietal gyrus thickness, lower PGSs for muscle weakness, and lower PGSs for reaction time and more active music engagement. Despite this, the meta-analyses and cohort-level analyses provide inconclusive evidence as to whether PGSs for neuromotor traits predict active music engagement.

The meta-analyses revealed that the PGS for walking pace was positively associated with active music engagement, with the strongest effect in CLSA. The genetics of walking pace are important for motor function and are connected to several domains of health. For example, faster self-reported walking is causally associated with lower cardiac and stroke risk (41,56) and genetically correlated with lower cardiometabolic, respiratory, psychiatric, and all-cause mortality risk and higher educational attainment (41,57). We observed patterns showing that genetic associations between walking pace and active music engagement may differ between males and females, which is expected, given the role of sex differences in brain development and aging (58–60). The meta-analyses across cohorts also revealed patterns of association between PGS for muscle weakness and active music engagement. Hand muscle weakness, measured by low hand grip strength, is a simple motor performance metric that provides a window into neural resources needed to commit a motor action, e.g., hand grip strength recruits corticospinal tract motor neurons (61). Hand grip strength also has genetic associations with frailty, cardiac, psychiatric, motor, and general health traits (62–66). Lastly, we also observed trends of associations with PGS for inferior parietal gyrus thickness and active music engagement. The inferior parietal lobe is critical for music processing because of its role in sensorimotor integration (Sun et al., 2022) and musical rhythm processing (68,69). Despite the potential relationship between PGS for inferior parietal gyrus thickness and music engagement, we did not observe significant meta-analyzed or cohort-level associations with other neuromotor traits.

Our results provide evidence for potential shared genetics between active music engagement motor traits, suggesting that investigations of “transfer” from musical to non-musical skills should consider individual differences. Several authors have suggested that genetic factors and gene-by-environment interactions likely confound the literature on the “transfer” to non-musical domains (3,3,24,70,71). Schellenberg and colleagues have evaluated the state of the literature on music-based “transfer” to non-musical domains of cognition, suggesting that music-induced benefits are weak correlations (3,71,72). Individual differences (not limited to genetic predispositions) possibly magnify the observed correlations between music engagement and other domains (71). Thus, gene-by-environment interactions can be incorporated into future music-based intervention studies to improve their rigour.

Our results suggest that PGSs for the rate of change in brain structure in motor regions do not account for differences in active musical engagement. However, Brouwer et al.’s age-independent meta-GWASs on the rate of change of brain structures capture the average rate of change across the lifespan rather than during specific developmental or aging periods. Moreover, our findings do not rule out the possibility that other plasticity traits may influence active music engagement. For example, the neural structure of white matter tracts, especially the right corticospinal tract organization, in infanthood predicts rhythmic and musical abilities in school-age children (73). Future investigations should also consider functional neuromotor phenotypes, e.g., the genetic architecture of functional brain networks (74), especially given the relevance of sensorimotor or audiomotor networks to active music engagement (6).

Although the meta-analyses revealed promising insights, we discuss some study limitations. While BioVU and OM had validated outcomes, the outcomes in WLS and CLSA had unknown psychometric profiles. The phenotypes also included singing together with musical instrument engagement, which may require different neuromotor resources, e.g. singing requires coordination of laryngeal, oropharyngeal, and facial muscles from the corticobulbar tract (75) and musical instruments require fine motor control of the upper limbs that descend from the corticospinal tract (2). Therefore, motor PGSs and active music engagement associations may be deflated due to this heterogeneity. Current initiatives, i.e., the *Musicality Genomics Consortium,* are tackling this issue by validating succinct measurements to implement in biobanks.

Second, the predictive power of PGSs are constrained by discovery and target cohort sample sizes and phenotype measurements (39), e.g., the childhood motor functioning and neuromotor discovery GWASs had smaller sample sizes. Additionally, current motor behaviour GWASs do not capture the full range of complex motor phenotypes, but recent studies show the feasibility of collecting motor learning data from thousands of individuals online (76) for future GWASs. Lastly, due to data availability, our primary analyses were constrained to genetic similarity with European ancestry due to differences in allele frequencies and the limited generalizability of PGSs across ancestries (77,78). Future work should leverage multi-ancestry methods and focus on musicality phenotyping in diverse populations.

## 5. Conclusion

PGS for higher self-reported walking pace may be associated with greater music engagement. However, analyses did not yield sufficient evidence to support that PGSs for neuromotor phenotypes predict active music engagement. Our results suggest that shared genetic factors for motor function may predict active music engagement, holding significance for longitudinal studies and interventions aiming to understand the transfer of musical learning to non-musical domains.

## Authorship Contributions

T.L.H.: Data curation, research design, data cleaning and quality control of genetic data, conducted all analyses, writing of first draft and writing of all sections on the manuscript; P.L.C.: Data curation (CLSA, WLS), data collection (OM), genetic quality control (OM), feedback on supplementary materials; D.E.G: Data collection (OM), feedback on study design for the pre-registration phase and feedback on the manuscript. Y.N.M.: Study design, data curation, and genetic quality control support (WLS, CLSA, OM); S.N.: Data collection (OM) and manuscript feedback; R.N.: Data collection (OM) and genetic quality control (WLS, CLSA, OM); A.C.S.: Data collection (OM), code review, and manuscript feedback; E.S.T.: genetic quality control support for CLSA and methods development for longitudinal summary statistics scoring; R.v.K.: methods development for longitudinal summary statistics scoring; D.F.: Supervision, data curation (CLSA), study design, statistical analyses feedback, methods development for longitudinal summary statistics scoring, manuscript feedback and revisions; M.H.T: funding, supervision, study design, manuscript feedback; R.L.G.: funding, supervision, data curation (all cohorts), study design, analyses feedback, manuscript feedback and revisions.

## Acknowledgements

This research was made possible using the data/biospecimens collected by the Canadian Longitudinal Study on Aging (CLSA). Funding for the Canadian Longitudinal Study on Aging (CLSA) is provided by the Government of Canada through the Canadian Institutes of Health Research (CIHR) under grant reference: LSA 94473 and the Canada Foundation for Innovation, as well as the following provinces, Newfoundland, Nova Scotia, Quebec, Ontario, Manitoba, Alberta, and British Columbia. This research has been conducted using the CLSA

Comprehensive Dataset version 7.0 and Genome-wide Genetic Data Release 3.0 under Application Number 2104030. The CLSA is led by Drs. Parminder Raina, Christina Wolfson and Susan Kirkland.

Vanderbilt University Medical Center’s BioVU projects are supported by numerous sources: institutional funding, private agencies, and federal grants. These include NIH-funded Shared Instrumentation Grant S10OD017985, S10RR025141, and S10OD025092; CTSA grants UL1TR002243, UL1TR000445, and UL1RR024975. Genomic data are also supported by investigator-led projects that include U01HG004798, R01NS032830, RC2GM092618, P50GM115305, U01HG006378, U19HL065962, R01HD074711, K07CA172294, 14GRNT20460090, P01DK038226, R24DK96527, U01HG004798, R01LM010685, R01NS032830, R01EY012118, K12HD043483, R01DK078616, RC2GM092618, APP1064524, R01CA162433, P01HL056693, P50GM115305, U01HG006378, U19HL065962, U01HG004603, P50CA09813, R01HD074711, R03HD078567, R01DK080007, and P50HL081009.

The National Institutes of Health supported research and data collection for the Vanderbilt’s Online Musicality study under award numbers R01DC016977, DP2HD098859, and UL1TR000445, as well as the National Science Foundation under award number NSF1926794. We thank all researchers and staff involved in Vanderbilt’s Online Musicality Study data collection.

T.L.H. is supported by the Data Science Institute at the University of Toronto, the School of Graduate Studies at the University of Toronto, CANSSI Ontario STAGE, and Mitac’s Globalink Research Award. Thank you to David and Marcia Beach for your summer study award, which supported the work for this project.

Thank you to the participants of all cohorts included in this study, to Dr. Daphne Tan for your mentorship throughout this project, and to Maria Niarchou for your support with the BioVU data. Thank you to the staff at the Wisconsin Longitudinal Study for your valuable support with the data resource.

## Data and Code Availability Statement

Individual data are available from the Canadian Longitudinal Study on Aging (www.clsa-elcv.ca) for researchers who meet the criteria for access to de-identified CLSA data. Individual data from the Wisconsin Longitudinal Study are available for researchers who meet the criteria to access de-identified data. Individual data from Vanderbilt’s BioVU repository cannot be shared publicly due to patient confidentiality; however, Vanderbilt University Medical Center researchers may request access (victr.vumc.org). Individual deidentified data for Vanderbilt’s Online Musicality will also be deposited to dbGaP. Study identifiers for all GWAS summary statistics used for polygenic score calculations are in Supplementary Table 2. Source summary level data for all figures and the output of all models are available in the supplementary materials. Code for all modelling and figures are available at: [insert osf link here].

## Disclaimers

The opinions expressed in this manuscript are the author’s own and do not reflect the views of the Canadian Longitudinal Study on Aging, BioVU, or the Wisconsin Longitudinal Study. The content is solely the authors’ responsibility and does not necessarily represent the official views of the National Institutes of Health. **The authors have no conflicts of interest to declare.**

## Notes

### Competing Interest Statement

The authors have declared no competing interest.

